# Recovery of high quality metagenome–assembled genomes from full–scale activated sludge microbial communities in a tropical climate using longitudinal metagenome sampling

**DOI:** 10.1101/2021.08.21.456929

**Authors:** Mindia A. S. Haryono, Ying Yu Law, Krithika Arumugam, Larry C.-W. Liew, Thi Quynh Ngoc Nguyen, Daniela I. Drautz-Moses, Stephan C. Schuster, Stefan Wuertz, Rohan B. H. Williams

## Abstract

Analysis of metagenome data based on the recovery of draft genomes (so called metagenome–assembled genomes, or MAG) have assumed an increasingly central role in microbiome research in recent years. Microbial communities underpinning the operation of wastewater treatment plants are particularly challenging targets for MAG analysis due to their high ecological complexity, and remain important, albeit understudied, microbial communities that play a key role in mediating interactions between human and natural ecosystems. In this paper, we consider strategies for recovery of MAG sequence from time series metagenome surveys of full–scale activated sludge microbial communities. We generate MAG catalogues from this set of data using several different strategies, including the use of multiple individual sample assemblies, two variations on multi–sample co–assembly and a recently published MAG recovery workflow using deep learning. We obtain a total of just under 9,100 draft genomes, which collapse to around 3,100 non–redundant genomic clusters. We examine the strengths and weaknesses of these approaches in relation to MAG yield and quality, showing the co-assembly offers clear advantages over single–sample assembly. Around 1000 MAGs were candidates for being considered high quality, based on single–copy marker gene occurrence statistics, however only 58 MAG formally meet the MIMAG criteria for being high quality draft genomes. These findings carry broader implications for performing genome–resolved metagenomics on highly complex communities, the design and implementation of genome recoverability strategies, MAG decontamination and the search for better binning methodology.

## Introduction

Over course of the last half decade the use of genome–resolved metagenome analysis has become a common approach for dealing with whole community metagenome data collected from microbiomes and complex microbial communities (Quince et al., 2017b). Starting with deeply sequenced genomic DNA, metagenome assembly is performed in order to reconstruct short fragments of the underlying member genomes, which are then analysed further using data clustering procedures (genome binning (Sangwan et al., 2016)) with the objective of recovering draft genomes of the member species, referred to as metagenome–assembled genomes (MAG). This approach, now readily deployable due to the availability of near–automated bioinformatics workflows (Uritskiy et al., 2018), has been successfully used on a great variety of microbial communities (Almeida et al., 2021; Nayfach et al., 2019; Parks et al., 2017; Pasolli et al., 2019; Singleton et al., 2021; Stewart et al., 2019; Tully et al., 2018) and has resulted in recovery of draft genomes for many new species that would have most likely remained uncharacterised due to a lack of knowledge of their required culture conditions (Parks et al., 2017).

Despite impressive accomplishments, the MAG approach still harbours many challenges and limitations. By nature, short read metagenome assemblies remain highly fractionated, resulting from the limited ability of short read sequencing to accurately capture complex repeat regions (Chen et al., 2020) and the difficulties encountered in reconstructing sequence from closely related strains or sub–species (Bertrand et al., 2019; Quince et al., 2020; Vicedomini et al., 2021; Quince et al., 2017a). In practice a draft genome obtained from these methods would contain at best, tens and, more typically, hundreds, of distinct contigs, and so there are inherent difficulties in accurately determining the degree of genome completeness and the extent of contamination from non-cognate genomes (Chen et al., 2020), and in identifying the presence of horizontally transferred sequence (Douglas and Langille, 2019). Another limitation relates to impact of the eco–genomic complexity of the community under study, both in terms of genomic diversity, particularly at sub–species or strain level, but also in terms of overall community richness and evenness (Quince et al., 2017b). When applied to microbial communities of high complexity, a typical MAG analysis will return many draft genomes of unremarkable quality, as defined by currently accepted criteria (Bowers et al., 2017). Some of these challenges may be addressed using emerging methods, such as longread sequencing (Arumugam et al., 2021; Singleton et al., 2021), synthetic long-read methods (Bishara et al., 2018) and adaptations of chromosome conformation capture methods (Bickhart et al., 2021; DeMaere and Darling, 2019). However all of these new techniques are themselves complex and will contain their own limitations, and since the vast majority of non–amplicon metagenome data has been collected using Illumina shotgun sequencing, there remains a clear need to develop more refined methods to recover genomes from short read metagenome assemblies.

Complex microbial communities associated with full–scale wastewater treatment plants (activated sludge) are particularly challenging targets for MAG–based analyses due to high species richness, high species evenness and extent of genetic diversity (Law et al., 2016; Pérez et al., 2019; Singleton et al., 2021; Yang et al., 2020; Ye et al., 2020). Recent comparative analyses undertaken with amplicon sequencing surveys suggest that these activated sludge communities are more complex than the host-associated microbiomes, including the human fecal microbiomes, by an order of magnitude (Wu et al., 2019). To date, several MAG-based analyses of activated sludge communities have been reported, varying in sequencing depth, raw sequence and availability of recovered genome (MAG) sequence, including one recently published study that employed long-read metagenomics (Singleton et al., 2021). In this paper, we consider strategies for recovery of MAG sequence from time series metagenome surveys of full–scale activated sludge microbial communities. We generate MAG catalogues from this set of data using several different strategies, including the use of multiple individual sample assemblies, two variations on multi–sample co–assembly and a recently published MAG recovery workflow using deep learning (Nissen et al., 2021). We examine the strengths and weaknesses of these approaches in relation to MAG yield and quality, and present a catalogue of non-redundant draft genomes comprised of at least putatively high quality under the MIMAG criteria. All raw data and high quality MAG sequence have been made available via NCBI (BioProject Accession PRJNA731554), and key data products, including metagenome assemblies and the complete set of recovered MAG sequence data, are being made publicly available on Zenodo (DOI 10.5281/zenodo.5215738).

## Results

### Summary of data obtained and overall study design

As part of a long–term sampling project surveying the microbial ecology of wastewater treatment in tropical climates, we sampled activated sludge from aerobic–stage tanks in a full-scale wastewater treatment plant in Singapore, known to perform enhanced biological phosphorus removal (EBPR) and previously studied by us in Law et al. (2016), obtaining 24 samples over approximately a 10 month period. The median sampling interval was 7 days (mean 13 days, with range 7-56 days). At each sampling event, we obtained samples for DNA extraction from the aerobic treatment tank (including a panel of co–assayed physico–chemical measurements), and performed whole community shotgun metagenome sequencing on all samples. In total, we obtained 1.5 billion reads with a mean of 62.6M reads per sample (range: 45.7M–101.4M; **Supplementary Table 1**). From these data we constructed catalogues of metagenome–assembled genomes (MAG) using several approaches as described below.

In our primary analysis, we performed both individual sample assembly of data from each of the 24 samples and co–assembly of the same ensemble of data (see **Methods: Genome–resolved metagenome analysis**), in order to formally compare the results of each of these two major approaches to MAG–based analysis. Metagenome assembly was performed using metaSPAdes (Nurk et al., 2017) and genome binning was performed using MetaBAT2 (Kang et al., 2019) in both cases. From the co-assembly, we generated two sets of MAGs, one using coverage profiles generated across all 24 samples and the other generated using the entire read set treated as a single meta–sample (see **Methods: Genome–resolved metagenome analysis**), which we refer to as multi–BAM and single–BAM co-assembly binning, respectively.

As a secondary analysis, we performed metagenome binning using a recently published deep learning workflow called VAMB (Nissen et al., 2021), which is described later in the article.

### Comparative analysis of individual sample assembly and co-assembly from MetaBAT2–based workflows

A total of 7,138 MAGS were recovered from the three types of assembly–binning workflows. Between 94 and 273 MAGs (mean 143) were obtained from each individual sample assembly, with a total of 3,429 MAGs being generated from all 24 individual assemblies (**Table 1** and **Supplementary Table 2**). Approximately 10% and 27% of individual sample assembly MAGs were candidates for being high (pHQ) and medium quality (MQ) under the MIMAG criteria (Bowers et al., 2017) (see **Methods: Genome quality estimates**). The single–BAM and multi–BAM co-assembly binning workflows returned 1,997 and 1,712 MAGs, respectively (**Table 1**). The proportions of pHQ– and MQ–MAGs obtained from co-assemblies were higher compared to those observed from the ensemble of individual sample assemblies (**Table 1**), with 14.3% and 17.7% being classifiable as pHQ–MAGs in the single–BAM and multi–BAM co-assembly binning, respectively, and approximately 30% of MAG from each type of co-assembly binning workflow, holding MQ status.

**Table 1:** Number of MAGs (percentage of the total observed within workflow) from different assembly–binning workflows categorised by initial quality evaluation. Percentage of total MAG number per workflow in brackets.

| Assembly-binning procedure | Total | Putative genome quality |  |  |  |
| --- | --- | --- | --- | --- | --- |
|  |  | High | Medium | Low | Unclassified |
| Individual assemblies ($n=24$) | 3429 | 341 (9.9%) | 934 (27.2%) | 1775 (51.8%) | 379 (11.1%) |
| Co-assembly, single-BAM | 1997 | 285 (14.3%) | 589 (29.5%) | 878 (44.0%) | 245 (12.3%) |
| Co-assembly, multi-BAM | 1712 | 303 (17.7%) | 532 (31.1%) | 641 (37.4%) | 236 (13.8%) |
| VAMB | 1941 | 156 (8.0%) | 475 (24.5%) | 1293 (66.6%) | 17 (0.9%) |

The proportion of reads mapped to co-assemblies was higher (mean 92%; *n*=2) than the proportion observed to map to individual sample assemblies (mean 67%, *n*=24) (**Supplementary Table 2**).

As expected, estimated MAG genome quality demonstrated a strong association with relative abundance–expressed as a normalised coverage measure that permits comparisons across workflows whose variable number of input sequence reads would bias estimation (see **Methods: Genome–resolved metagenome analysis**)–with the proportion of pHQ MAGs being highest in the top 10%-ile of the normalised coverage, and decreasing in a roughly uniform manner thereafter (**Fig. 1**). A similar trend was observed for MQ–MAGs, and the proportion of poor quality MAGs expanded in the bottom 50% of the normalised coverage distribution.

**Figure 1:**
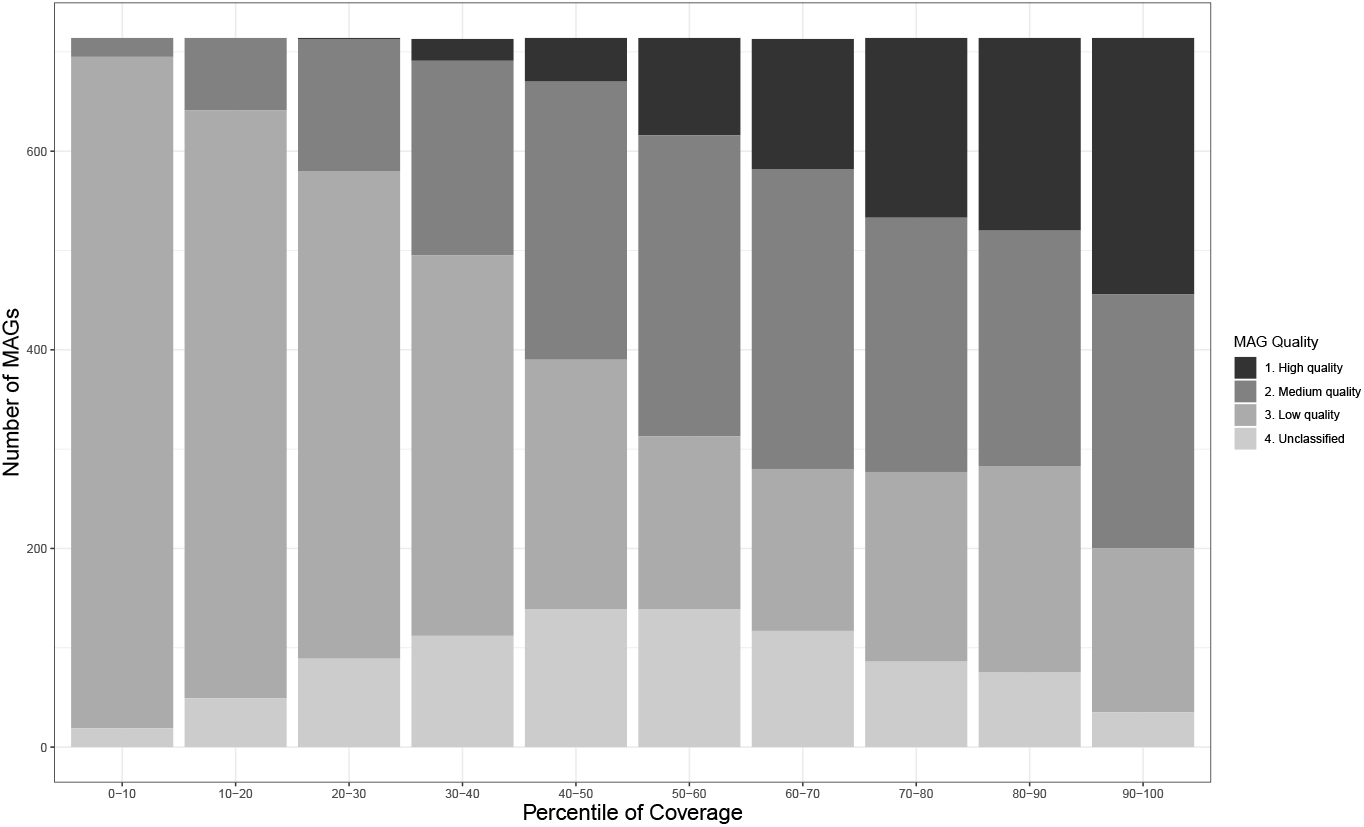
Relationship between genome quality and relative abundance within the MetaBAT2–derived MAG catalogues obtained in this study. Relative abundance is inferred using a normalised measure of coverage, permitting comparison of mean MAG coverage among the three different assembly–binning workflows (see Table 1 and Methods: Genome recovery from individual sample assemblies). Within each decile of the normalised coverage distribution, the numbers of MAG meeting each of the four MIMAG– derived quality levels is shown.

Given the expected high degree of genomic redundancy among the complete set of 7,138 MAGs generated from the three assembly–binning workflows employed, the entire set was de-replicated and grouped into non-redundant genome clusters (*secondary clusters* as defined by the dRep workflow (Olm et al., 2017); see **Methods: Genome de–replication procedures**). In total 2,912 non–redundant clusters where obtained, comprised of between 1 and 26 MAGs (median 2; mean 2.45) (**Supplementary Table 3**). Of these 2,912 secondary clusters, 382 (13.1%) contained at least one MAG that was pHQ, and 690 (23.7%) contained MAGs that were MQ at best, with the remainder containing MAGs of either low quality (LQ; *n* =1576; 54.1%) or else unclassifiable (UC; *n* =264; 9.1%).

To gain further insight into the effectiveness and inter–relationship of each type of genome recovery workflow, the set of 2,912 non–redundant clusters were further categorised according to the the types of workflow which had contributed at least one genome to a given non-redundant cluster (**Supplementary Table 4** and **Fig. 2A**). Of these 2912 secondary clusters, 346 (11.9%) contained genomes contributed from both individual sample assemblies and both types of co–assembly binning procedure, and 1070 (36.8%) contained MAGs recovered from type of co–assembly but not any individual sample assembly (**Fig. 2A**). Relatively few MAGs were observed arising from individual sample assembly and from either, but not both, types of co-assembly (23 and 14 secondary clusters, respectively, against single–BAM and multi-BAM; **Fig. 2A**). In contrast, we observed substantial numbers of secondary clusters that were only comprised of MAGs obtained from within one of the three workflows (**Fig. 2A**).

**Figure 2:**
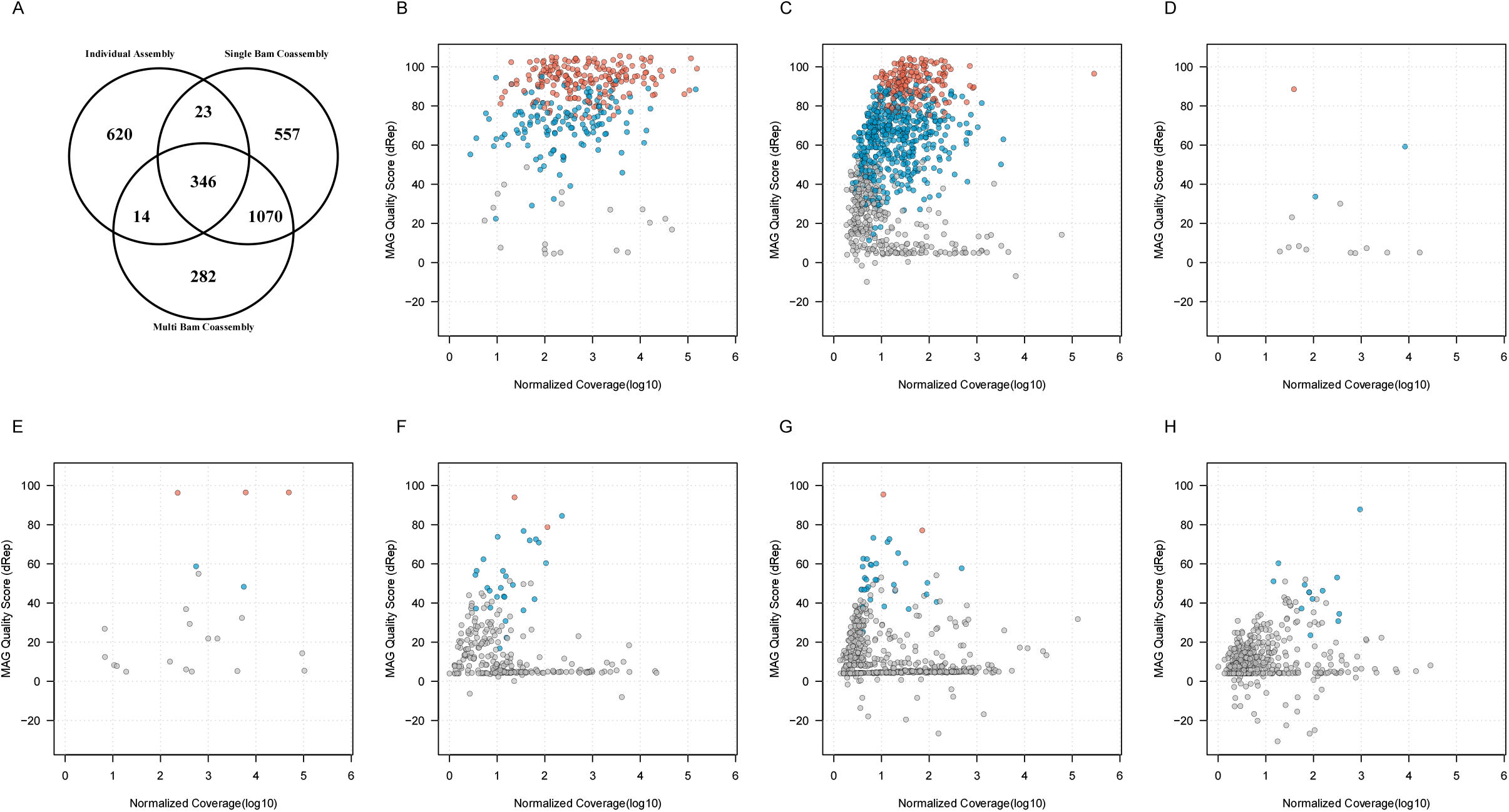
Inter–relationships between genome quality, relative abundance and recovery workflow type for 7.138 MAGs recovered from the use of MetaBAT2. *A*) secondary cluster membership by workflow (7,138 MAG categorised into 2,912 secondary clusters; the remaining panels show the highest observed genome quality with a secondary cluster (vertical axis) against normalised coverage measure (horizontal axis) for secondary clusters *B*) containing MAGs from all three workflows (*n*=346); *C*) recovered from both single–BAM and multi–BAM co–assembly workflows (*n*=1070); *D*) recovered from both individual assemblies and multi–BAM co-assembly (*n*=14); *E*) recovered from both individual assemblies and single–BAM co-assembly (*n* =23); *F–H*) recovered solely from multi–BAM co-assembly (*n* =282), single–BAM co-assembly (*n* =557) and individual assemblies (*n* =620), respectively. Secondary clusters containing pHQ, MQ, and LQ MAGs are coloured in red, blue, and grey, respectively.

We then examined how these associations were patterned by genome quality and relative abundance, using a composite quality statistic as defined in the dRep pipeline and a normalised measure of MAG coverage that adjusted for differences in coverage that are present across the three types of workflows (**Fig. 2B–2H**). Each secondary clusters was represented by the best quality MAG observed in that cluster, as defined by the maximum dRep quality score within the highest MAG quality category from that cluster.

We observed that the 346 secondary clusters comprised of MAGs recovered from all three workflows had the highest overall coverage and over half of these secondary clusters contained at least one pHQ genome (189/346 or 54.6%). In the larger set of 1070 secondary clusters that arose from both types of co-assembly workflow, 185 (17.3%) and 483 (45.1%) of these held at least one genome of pHQ and MQ level, respectively. These secondary clusters were also distributed across a lower coverage range than the previous category (**Fig. 2G**), consistent with the expectation that co–assembly procedures can recover genomes of less common or rare taxa. Of the remaining set of 1496 secondary clusters from the remaining four categories there were only 8 (0.54%) which held candidates for being pHQ–MAGs, with the remainder holding unremarkable or frankly poor quality (**Fig. 2B–2F**).

We then undertook several secondary analyses to examine whether co-assembly or individual sample assembly showed any inherent biases in genome quality (**Fig. 3**). Firstly, for secondary clusters that contained MAGs from all three workflows (**Fig. 2A–2B**) we examined the proportion of pHQ–MAG in secondary cluster that came from either type of co-assembly or from an individual assembly, but observed no clear pattern in relation to the origin of pHQ–MAGS (**Fig. 3A**). Secondly, we compared completeness and contamination statistics within a subset of 48 secondary clusters that contained at least one pHQ genome sourced from co-assembly and at least one pHQ genome from an individual assembly. Removing all genomes that did not attain pHQ status, on average this subset of secondary clusters contained 1.8 pHQ–MAGs (range: 1–2) sourced from the co–assembly workflows and 6.3 pHQ-MAGs (range: 1–22) arising from the individual assembly workflow. We calculated median completeness and median contamination within each secondary cluster, conditioned on workflow type, observing a bias towards higher completeness (**Fig. 3B, 3D**) and a lower contamination (**Fig. 3C, 3D**) in co–assembled genomes relative to genomes obtained from individual assemblies. Collectively, these data suggest that if we focus attention on recovered genomes that are plausibly of high quality, then these results indicate, in the communities studied here, that co-assembly conveys clear advantages in regards MAG yield.

**Figure 3:**
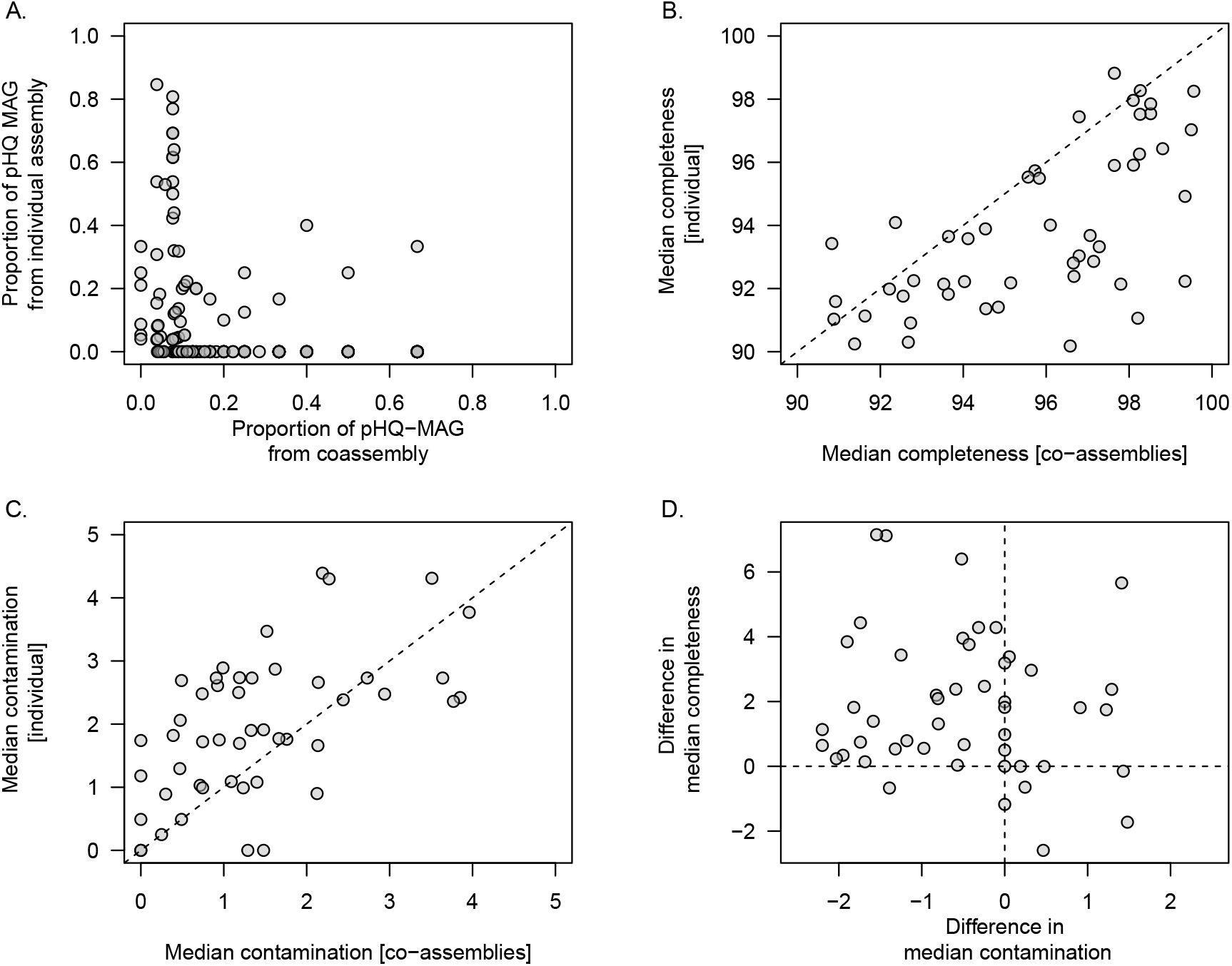
Examination of genome quality in MAGs recovered among different MetaBAT2-derived workflows. Each data point references a secondary cluster of highly related genomes obtained from the dRep workflow. *A*) proportion of MAGs within a secondary cluster that attain pHQ quality status from co–assembly workflows (*x*–axis) and individual assemblies (*y*–axis); *B–D* comparison of genome quality measures from within 48 secondary clusters in which at least one pHQ MAG is observed from a co-assembly workflow and from an individual assembly; *B*) median completeness (*C*_*p*_) observed from co–assemblies (*x*–axis) and individual assemblies (*y*–axis); *C*) median contamination (*C*_*n*_) observed from co–assemblies (*x*–axis) and individual assemblies (*y*–axis); *D*) relationship between the difference in median completeness observed in co–assemblies with respect to individual assemblies (*y*–axis) and difference in median contamination (co– assemblies with respect to individual assemblies) (*x*–axis).

### Decontamination of recovered draft genomes

To improve the number of high quality MAGs produced from the workflows above, we applied RefineM (Parks et al., 2017) to all MAGs from the three assembly procedures that possessed completeness of more than 90% (1,307 MAGs, contributed from 550 secondary clusters) regardless of their contamination and strain heterogeneity levels, as calculated by CheckM (Parks et al., 2015), considering these suitable candidates for refinement to high quality.

The results of the decontamination analysis are summarised in **Table 2A**. Of the 1,307 MAGs, 929 (71.1%) were classified as pHQ prior to decontamination, and of these, 855 (92.0%) retained the same quality level after application of RefineM. Of the remaining 74 pHQ–MAGs, the majority 94.6% were converted to MQ, with only four being reduced to LQ status. Of the 127 MAGs originally classified as MQ, 35.4% (45/127) attained pHQ status following application of RefineM. In the set of 251 MAGs that held UC status before decontamination, only 7.2% and 16.7% improved their quality to pHQ and MQ, respectively, suggesting that most MAGs that hold contamination above 10% are likely to be of highly flawed construction.

**Table 2:**
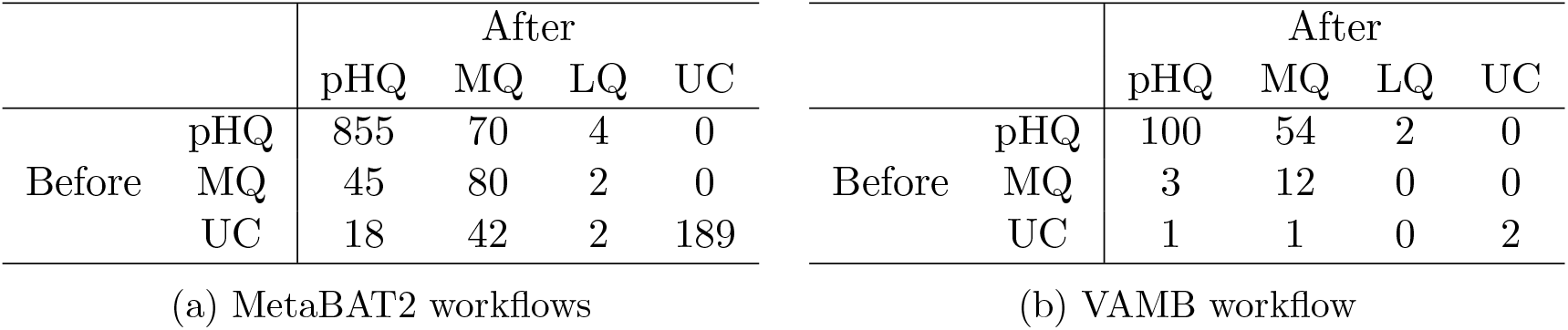
Number of MAGs categorised by genome quality assignments before and after decontamination with RefineM for (a) MAGs obtained from MetaBAT2 workflows and (b) MAGs from VAMB workflow. Input MAGs held CheckM–estimated completeness*>* 90%.

Across all MAGs, the average number of contigs removed by RefineM was 59 (range: 0–3,309) with CheckM completeness, contamination, and strain heterogeneity reduced on average by 1.7%, 5.8%, and 0.5, respectively (**Supplementary Table 5**).

### Genomes recovered using a deep variational autoencoder workflow (VAMB)

As a secondary, complementary analysis to the canonical approach taken above, we performed genome recovery using a recently described workflow called VAMB that utilises deep variational autoencoders (Nissen et al., 2021). Using data from the 24 individual– sample assemblies, VAMB generated 1,941 MAGs of minimum total sequence length of 200kbp (to match that used by the default MetaBAT2–based workflows).

Of the recovered draft genomes, 8.0%, 24.5%, 66.6%, and 0.9% were classified as pHQ, MQ, LQ and UC, respectively (**Table 1** and **Supplementary Table 6**). The pHQ–MAGs from VAMB were strongly associated with those detected by the MetaBAT2 workflows, with only 1 and 5 secondary clusters containing pHQ–MAGs (**Figure 4A**) and MQ–MAGs (**Figure 4B**), respectively, that were not recovered by any other workflow. While a substantive number of secondary clusters containing LQ–MAGs were recovered by VAMB (**Table 1** and **Figure 4C**), interestingly, the number of secondary clusters containing UC–MAGs was two orders of magnitude lower than the number observed in the MetaBAT2 workflows (**Figure 4D**). This suggests that the VAMB methodology may likely provide superior control of gross contamination, although possibly at the expense of recovery of more complete, higher quality MAGs.

**Figure 4:**
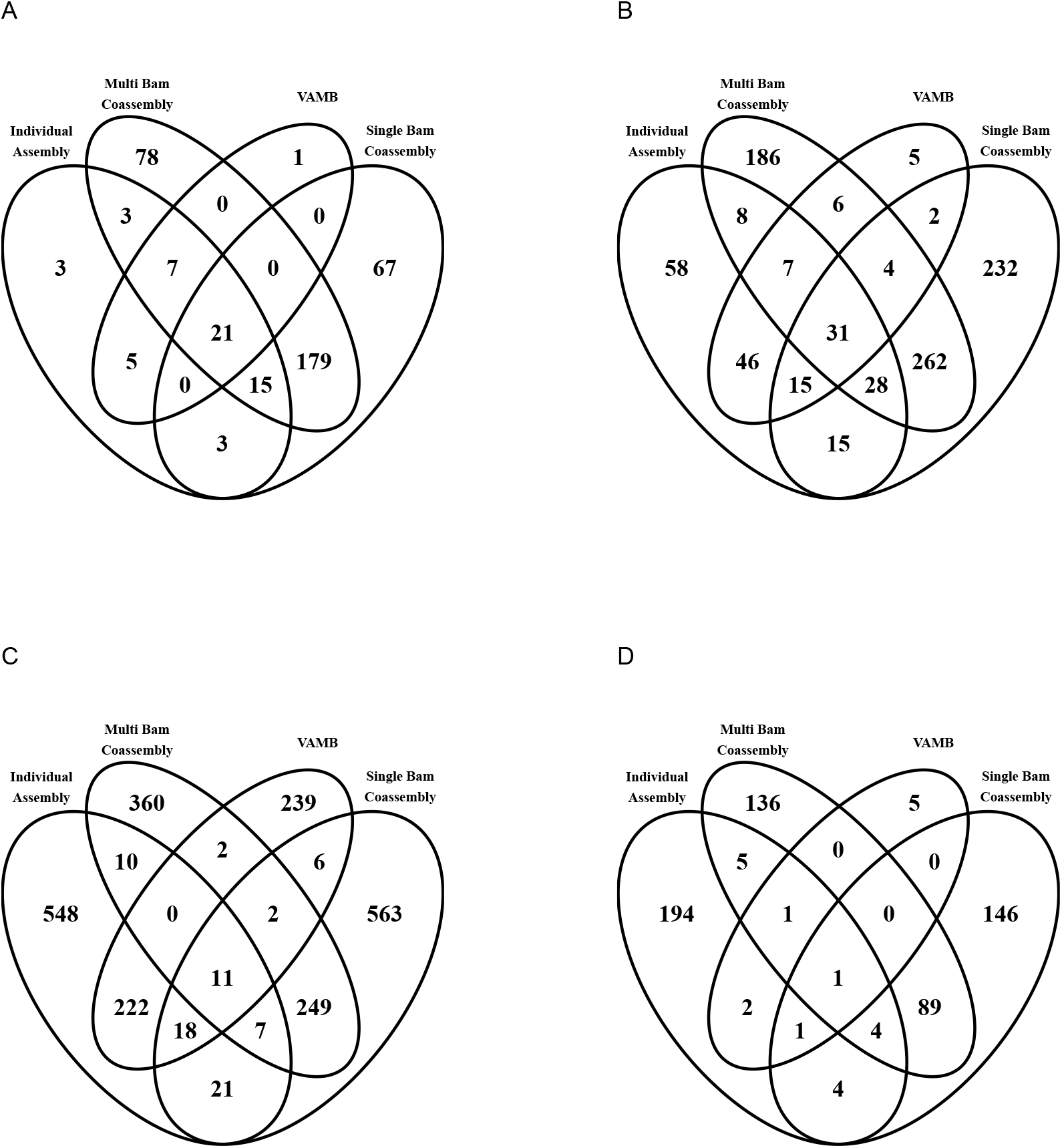
MAG recovery across all four workflows employed in this study, conditioned by genome quality. The entire set of 9,709 MAGs was categorised by genome quality level, and within each quality genome level further categorised by the workflow of origin observed with each secondary cluster. Venn diagrams showing interrelationships are shown here for *A*) pHQ (putative high quality); *B*) MQ (medium quality); *C*) LQ (low quality) and *D*) UC (unclassified).

As above, we applied RefineM workflow to the 175 MAGs with completeness above 90% (contributed from 36 distinct secondary clusters), which were primarily comprised of pHQ–MAGs (*n* =156, 89.14%). After application of the RefineM workflow, 59% of pHQ-MAGs retained their quality status, and 38% were reduced to MQ–status. The numbers of MAGs in remaining categories was low (**Table 2B**). The average number of contigs removed by RefineM was 53 (range: 0-495), and completeness, contamination and strain heterogeneity were reduced on average by 4.6%, 1.4% and 1.3 units, respectively (**Supplementary Table 6**).

### Catalogue of high quality genomes from tropical climate activated sludge

The entire set of MAGs recovered from all four sources were combined into a single set of 9,079 bins (7138 bins from the MetaBAT2 workflows and 1941 from VAMB; see **Supplementary Table 7**), corresponding to 3,113 secondary clusters, as defined by dRep. Of the 9079 MAGs defined by this analysis, 1085 (11.9%) were categorised as pHQ. Of these 1044 (96.2%), 124 and 5 were comprised of less than 500, 50 and 10 contigs or less, 1066 MAGs (98.3%) held an N50 of at least 10kb and 142 MAGs (13.1%) contained at least one copy of the 5S, 16S and 23S SSU–rRNA genes. The 1085 pHQ MAGs were split among 382 different secondary clusters.

Taxonomic analysis (**Fig. 4**) showed a predominance of phyla *Bacteroidota* and *Proteobacteria*, which accounted for 44.4% (482/1085 MAGs within 100 secondary clusters) and 20.6% (223/1085 MAGs in 98 secondary clusters) of MAGs classified as pHQ. Other phyla that were observed at relative frequencies above 1% were *Chloroflexota* (5.4%, 59 MAGs within 22 secondary clusters), *Planctomycetota* (5.3%, 57 MAGs within 37 secondary clusters), *Spirochaetota* (4.4%, 48 MAGs within 7 secondary clusters), *Actinobacteriota* (3.8%, 41 MAGs within 19 secondary clusters), *Acidobacteriota* (2.7%, 29 MAGs within 17 secondary clusters), *Myxococcota* (2.3%, 25 MAGs within 17 secondary clusters), *Nitrospirota* (2.3%, 25 MAGs within 4 secondary clusters), *Bdellovibrionota* (2.7%, 29 MAGs within 18 secondary clusters) and *Verrucomicrobiota* (2.3%, 25 MAGs within 16 secondary clusters).

Across the 382 secondary clusters, only 14 (3.7%) were comprised of MAGs annotated to species level and 155 (40.6%) were annotated to genus level, highlighting that over half the recovered pHQ MAGs were likely to be previously uncharacterised. Species-level annotations were observed for the polyphosphate accumulating organisms (PAO) *Candidatus* Accumulibacter SK–02 (*n* =2 MAGs) (Skennerton et al., 2015) and the cyanobacteria *Obscuribacter phosphatis* (Soo et al., 2014; Stokholm-Bjerregaard et al., 2017) (*n* =2 MAGs), and the glycogen accumulating organism, *Candidatus* Competibacter (McIlroy et al., 2014) (*n* = 1 MAG). Interestingly, we recovered a single MAG from *Romboutsia timonensis*, a member of the human gut microbiome (Ricaboni et al., 2016), and to our knowledge not previously identified in activated sludge communities, and genomes of the methane–oxidising bacteria *Methylosarcina fibrata* (Hamilton et al., 2015) (*n*=2 MAG). Genomes from the denitrifier *Hyphomicrobium denitrificans* (Martineau et al., 2014) were recovered (*n*=2 MAG), along with genomes from two species within the UBA2359 lineage within order *Chitinophagales* (GTDB), namely *Sphingobacteriales* bacterium TSM CSS and *Sphingobacteriales* bacterium TSM CSM and genomes from recently identified novel lineages in phyla *Bacteroidetes, Chloroflexi* and *Chlorobi* (see **Supplementary Table 8** for full details of species-level identifications).

The ammonia–oxidising bacteria (AOB), *Nitrosomonas* (Kowalchuk and Stephen, 2001) (*n*=16 MAG), and the nitrite oxidising bacteria (NOB), *Nitrospira* (Vijayan et al., 2021) (*n*=25 MAG), both key functional species in activated sludge-mediated bioprocesses, where only represented at genus level.

When we applied the more stringent form of the MIMAG criteria for high quality genomes (Bowers et al., 2017), that is, those with at least 18 tRNAs and the presence of a complete set of rRNA genes, only 58 HQ-MAGs were identified. In addition, 6 more high quality MAGs were recovered from RefineM pipeline, resulting in a total 64 high quality MAGs submitted to NCBI.

### Comparison to other MAG catalogues recovered from activated sludge

We systematically compared our MAG catalogue to several others that have been recently obtained (Singleton et al., 2021; Ye et al., 2020), using genome de–replication (see Methods: Genome de–replication procedures) and same criteria recently used in a comparative analysis of MAG catalogues from multiple cow rumen microbiomes (Watson, 2021). Collectively, this analysis defined a total of 6,328 secondary clusters, containing on average 1.9 MAGs (median 1.0; range 1–54 MAG). We examined the membership of these secondary clusters in relation to catalogue of origin (**Fig. 5**). Only 7 secondary clusters contained genomes from all three source catalogues. A larger proportion of related genomes (*n* =314) was observed between our catalogue and that of Ye et al. (2020), than between our study and the catalogue of Singleton et al. (2021), which may reflect the more diverse geographies and mixture of operational regimes incorporated in the former study. We highlight however, that our analysis is retrospective and thus should be interpreted with caution.

**Figure 5:**
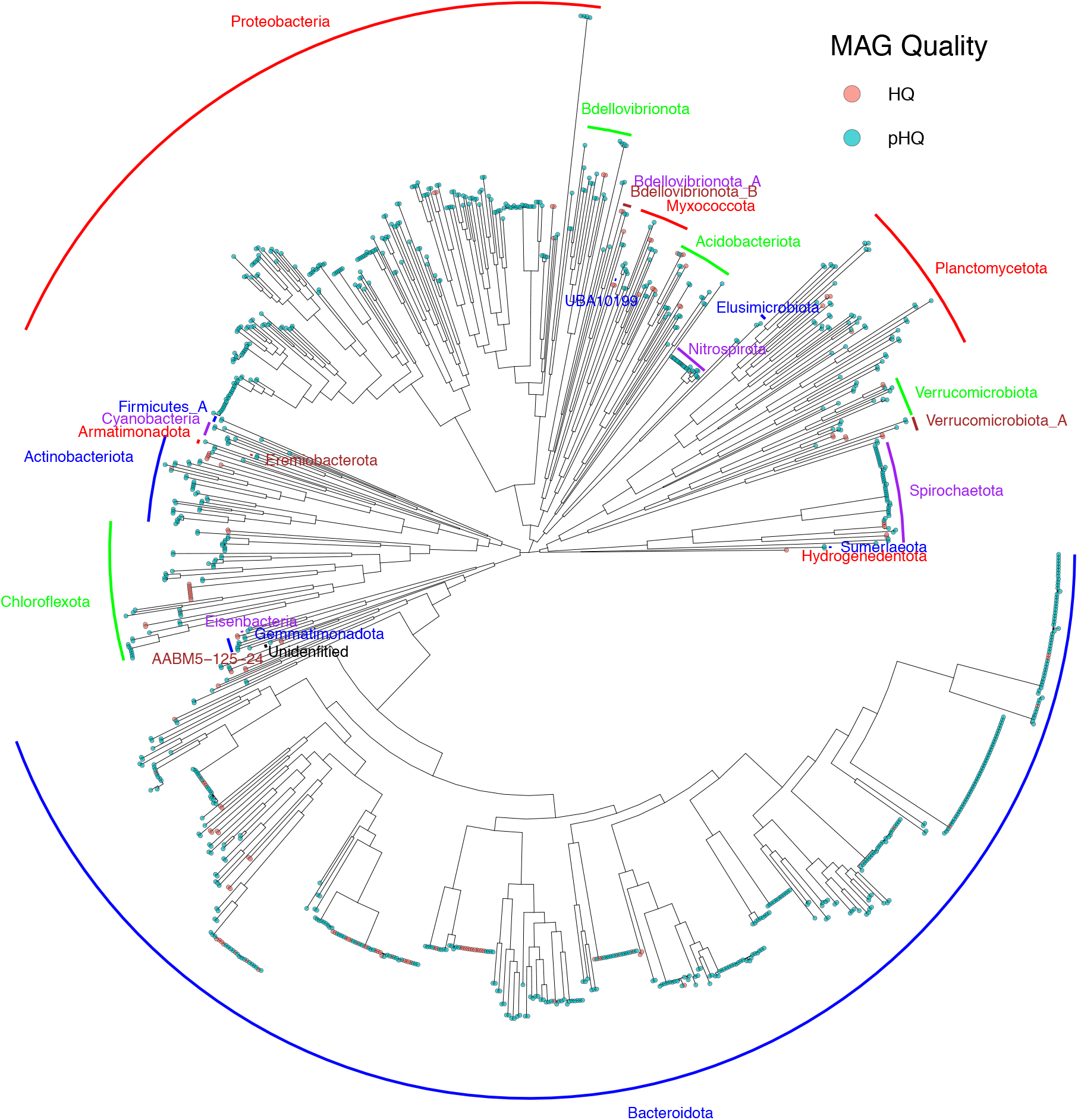
Phylogram of pHQ MAGs recovered from all four workflows used in this study. Phylum level annotations are listed as text labels. MAG holding pHQ status are highlighted in blue–green and the subset of those that hold HQ status under the MIMAG are highlighted in light red.

**Figure 6:**
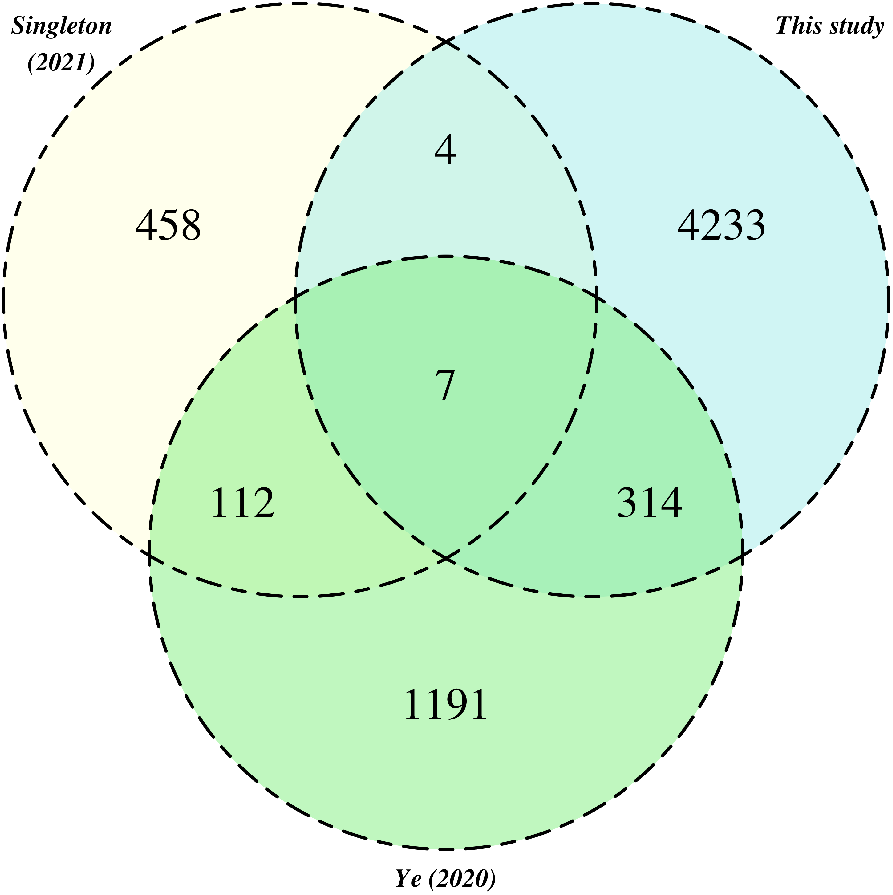
Relationships between pHQ-MAGs recovered in this study and MAGs from extant activated sludge catalogues. Counts reference secondary clusters categorised by the presence of MAGs originating from one of the three MAG catalogues.

## Discussion

In this paper we undertake a comprehensive genome–resolved metagenome survey of an activated sludge microbial community from a full–scale, tropical climate wastewater treatment plant, based on a time–series survey design. We obtain a total of just under 9,100 draft genomes, which collapse to around 3,100 non–redundant genomic clusters (defined under a stringent degree of relatedness). Around 1000 MAGs were candidates for being considered high quality, based on single–copy marker statistics (referred to pHQ in our analysis) but 58 MAGs formally meet the MIMAG criteria for being high quality draft genomes. In building these MAG catalogues, we undertake a systematic comparison of MAG recovery strategies, based on the use of individual–sample assemblies and two variations on the use of co–assemblies (using the combination of metaSPAdes for performing assemblies and MetaBAT2 for genome recovery). Additionally, we compared these results to those obtained from the use of a recently released deep learning variational autoencoder called VAMB (Nissen et al., 2021), which appears to convey some advantages in relation to control of MAG contamination. As discussed below, these findings carry broader implications for conducting genome–resolved metagenomics on highly complex communities.

The genomes recovered at pHQ level in this study represented 11 phyla, captured at a relative frequency of above 1%, with just under half being members of *Bacteriodota* and *Proteobacteria*, and represent the most comprehensive catalogue obtained from tropical climate activated sludge communities, building on our previous efforts (Arumugam et al., 2019, 2021; Law et al., 2016; Qiu et al., 2020). Wastewater microbial communities from tropical climates are understudied relative to their temperate climate counterparts, as are the bioprocesses that they support. Given the urgent need to understand the impact of climate change at microbial scales of life (Cavicchioli et al., 2019), such communities will become an increasingly important target of study, given their role as mediators of the interface between human and natural ecosystems (McLellan et al., 2015). In the present case, we obtained pHQ or confirmed HQ MAGs for expected taxa conveying key functionality to activated sludge bioprocesses including the AOB *Nitrosomonas*, NOB *Nitrospira*, the PAO *Candidatus* Accumulibacter and the GAO *Candidatus* Competibacter. Notable unexpected findings included, but were not limited to, the cyanobacterial PAO species *Obscuribacter phosphatis* and *Romboutsia timonensis*, previously identified in the human gut and plausibly an immigrant species from that source.

The large proportion of recovered genomes that hold unremarkable quality is expected, given the recognised challenges of performing these analysis on highly complex microbial communities (Pasolli et al., 2019) and the known complexity of full–scale activated sludge microbial communities, which are estimated to be more complex than the human gut microbiome by around an order of magnitude (Wu et al., 2019). The complexity of the community in regards to taxonomic novelty is also seen in the fact that around 60% of the recovered pHQ hold no assignment at species or genus level, and by the relatively low degree of recapitulation of genomes from other activated sludge catalogues. Consistent with the recognised limitations of MAG analyses conducted from short-read sequence data, the recovered genomes are unlikely to be resolved to strain level, and the size and complexity of the dataset limited the use of a recent genome–bin workflow for strain deconvolution (Quince et al., 2020) (data not shown). Nonetheless, the kind of densely sampled longitudinal data collected here is ideally suited for developing such strain–aware genome recovery methods.

As part of this analysis, we have undertaken a comprehensive comparison of individual sample assembly and co-assembly approaches for genome recovery, which has been relatively unexplored in the literature to date. Current thinking on MAG analysis suggests that assembling data from individual samples will aid the recovery of higher quality, relatively abundant genomes, while co-assembly will assist in the recovery of lower abundance genomes with the trade-off of artefacts associated with multi–sample analysis (Hofmeyr et al., 2020; Pasolli et al., 2019), including cross-sample chimeras (Chen et al., 2020), split–bins (Arumugam et al., 2021) and increased probability of recovering pan–genomic level (Chen et al., 2020), although this will no doubt be dependent on, the nature of the co-assembled samples (longitudinal versus cross-sectional; (Pasolli et al., 2019)), sample replication number, genetic diversity, community complexity and, of course, sequencing depth. In their comparative analysis on co–assembly and individual assembly of infant and maternal gut microbiomes, Pasolli *et al*. found little difference in number or quality of recovered genomes from either method, which included an analysis of both longitudinal and cross-sectional sampling designs, concluding that application to longer time–series would likely result in higher MAG yields (Pasolli et al., 2019). The findings of the present study are consistent with that view, with substantially higher numbers of pHQ–level MAGs being recovered from co-assembly procedures. We find some clear indications that co-assembly is advantageous in regards to genome quality, and, at least in the subset of MAGs that are recovered at pHQ level by both approaches, there is clear evidence that co-assembly will provide cognate MAGs with higher completeness and lower contamination statistics, as defined by single copy marker gene analysis: the extent to which this is generalisable to other settings is unclear.

Unexpectedly, we find the two specific modes of co–assembly are each capable of high MAG yields suggesting that greater depth *per se*, as implemented in the single–BAM approach, will recover almost as many pHQ MAGs (285 versus 303; **Table 1**) as the canonical differential coverage approach (multi–BAM). Additionally, the computational overheads of co–assembly can be substantial, as seen in the present case, and which may be untenable or impractical in some settings. Obviously this would also influence the choice of metagenome assembler, for example, MEGAHIT may be a more suitable choice of assembler than metaSPAdes for datasets at, or above, the scale of data employed here. Interestingly, as applied to this dataset, the deep–learning based VAMB workflow recovered pHQ MAGs that largely recapitulated those from the MetaBAT2 workflows. Collectively, these findings reinforce the view that MAG recovery is highly context-specific in relation to the community under study (Vollmers et al., 2017).

There remains an urgent need for methods to identify non–cognate contigs in fractionated assemblies, with the impact of contamination on gene–level becoming more widely recognised (Arkhipova, 2020), and one recently published analysis suggests that up to 15–30% of publicly–available MAGs classified at pHQ level will harbour chimeric content (Orakov et al., 2021). In the present study, we have examined removal of possible contamination using the RefineM workflow (Parks et al., 2017). Our results shed light on the strengths and weaknesses of the different recovery workflows we employed. From the MetaBAT2 workflows, there was a high degree of robustness in the case of recovered genomes that were classified at pHQ level, with over 90% retaining their pHQ status upon the application of RefineM. In the case of draft genomes that held high levels of contamination upon a backbone of high completeness, most also remained within the same genome quality category following de–contamination, suggesting that these recovered sequence constructions are fundamentally flawed. In the case of the bins recovered from VAMB, while around one third of pHQ changed quality level to MQ, there was an under–representation of complete genomes initially showing high degrees of contamination, suggesting that VAMB may be quite robust to the formation of chimeras. Whether this is a general property, or a consequence of the high redundant nature of time-series data, is a subject for further study.

Collectively, our results reinforce the ongoing need for analysis procedures suitable for recovering high quality MAGs from metagenome data, also highlighted by recent calls for more careful manual curation of recovered genomes (Chen et al., 2020; Lui et al., 2021) and the use of complementary sequencing, including long read (Arumugam et al., 2021; Singleton et al., 2021), synthetic long read (Bishara et al., 2018) and chromosome confirmation capture methods (Bickhart et al., 2021; DeMaere and Darling, 2019). Another relevant development is the direct use of assembly graphs in MAG recovery, including for the recovery of strain level sequence (Brown et al., 2020; Mallawaarachchi et al., 2020; Quince et al., 2020). Further attention could also be placed on the use of alternative feature representations for contig sequence and/or coverage data: most methods developed to date have used Euclidean space (of various dimensionality, ranging from two to several hundred), but other representations may hold substantive advantages, for example hyperspherical or hyperbolic embeddings (Ding and Regev, 2021) or related manifold learning methods *e*.*g*. as implicit in the use of VAMB (Nissen et al., 2021).

## Methods

### Metagenome extraction and sequencing

The field sampling methodology, sample handling, DNA extraction and sequencing methods have been previously described by us (Law et al., 2016). At a full-scale operational wastewater treatment plant in Singapore, treating mostly waste of domestic origin, we sampled the aerobic stage of an activated sludge tank known to perform enhanced biological phosphate removal (EBPR). At each sampling event, we obtained multiple samples for DNA extraction from the aerobic treatment tank and collected a panel of relevant physico–chemical measurements (data not analysed in this paper). Samples were snap frozen in a liquid nitrogen dry shipper immediately upon retrieval from the tank, and transported to the laboratory for subsequent genomic DNA extraction and sequencing on Illumina HiSeq2500 using a read length of 251bp (paired end) (see (Law et al., 2016)for details of all gDNA extraction, library preparation and sequencing protocols).

### Genome–resolved metagenome analysis

Unless otherwise stated data analysis was performed in the R Statistical Computing Environment (version 4.0.5) (R Core Team, 2021).

### Initial data processing

The raw FASTQ files were processed using cutadapt (version 1.5, with default arguments except `--overlap 10 -m 30 -q 20`) (Martin, 2011).

### Genome recovery from individual sample assemblies

From the processed read data, we initially performed individual sample assemblies using SPAdes (Nurk et al., 2017) in –meta mode with maximum *k*–mer value of 127, and performed metagenome binning using MetaBAT2 version 2.12.1 (default settings) to obtain an initial set of MAGs from each sample. Coverage for each contigs were extracted from SPAdes *k*–mer coverage, converted to log scale, and averaged per bin. To compare estimates of MAG coverage between samples, we normalised MAG coverages by centering using the per–sample mean per-MAG coverage, scaling by the per–sample standard deviation of coverage and then placing back on a positive scale by subtracting the smallest normalised coverage value across the entire set of MAGs.

### Genome recovery from co-assemblies

Processed read data from the 24 samples were co–assembled with SPAdes–3.13.0 (Nurk et al., 2017) (default parameters except `--meta -m 2900 -k 21,33,55,77,99,127 -t 50`). Binning was performed on contigs over 2500 bp in length with MetaBAT2 (Kang et al., 2019) (version 2.12.1 with default parameters except `-d -t 40 -m 2500 -v`), employing two different approaches, namely: *1*) using contigs from the co-assembly and 24 sorted .bam files made by aligning reads from each of the 24 datasets to the contigs from co-assembly, referred to as the *multi–BAM co–assembly* and *2*) using contigs from the co–assembly and a single sorted bam file made by aligning all reads from the 24 datasets to the contigs from co-assembly (referred to as a *single–BAM co–assembly*).

### Genome recovery using deep variational autoencoder (VAMB)

The recently published metagenome binner VAMB was employed on the 24 individual sample assemblies, following the described procedure in (Nissen et al., 2021). Briefly, all assembled contigs with minimum length of 2.500bp were compiled into a FASTA catalogue (`-m 2500 --nozip`). Processed read data from each of the 24 samples were mapped to this catalogue using Bowtie2 (version 2.3.4.3) (Langmead and Salzberg, 2012) and Samtools (version 1.9) (Li et al., 2009), with read depth being calculated using the MetaBAT2 script `jgi summarize bam contig depths`; default settings). We then ran VAMB (version 3.0.2) on the catalogue and read depth data using default parameters except for minimum total sequence length set at 200kb (`-o C` as the sample separator and `--minfasta 200000`).

### Genome quality estimates

Genome quality estimation of all all bins obtained from all four different pipelines (individual sample assemblies, single–BAM co-assembly, multi–BAM co-assembly and VAMB) was performed by running the CheckM (version 1.0.13) (Parks et al., 2015) `lineage wf` workflow using default parameters (except `-t 20 -x fa` or `-t 20 -x fna` for VAMB bins). The output was then tabulated with the CheckM `qa` command using 20 threads (`-t 20`). MAG quality was then classified using the MIMAG criteria (Bowers et al., 2017) with modifications as follows: *1*) MAGs with CheckM completeness (*C*_*p*_) and CheckM contamination (*C*_*n*_) values *>*90 and *<*5, respectively, were classified as candidates for being high quality (pHQ) genomes bins; *2*) MAGs with *C*_*p*_ *>*= 50 and *C*_*n*_ *<* 10 were categorized as being of putatively medium quality (MQ) ; *3*) MAGs with *C*_*p*_ *<* 50 and *C*_*n*_ *<* 10 were classified as candidate low quality (LQ) and *4*) MAGs that did not fall into any of the above three categories were unclassified (UC). The N50 value (*N*_50_) for each MAG was calculated using QUAST version 5.0.0 (Gurevich et al., 2013) with flags `--mgm --rna-finding --min-contig 1 --max-ref-number 0`. for each MAG, we computed an overall (univariate) quality statistic, *Q*_*d*_ as defined by within the dRep workflow (Olm et al., 2017), defined as 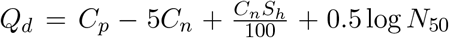. MAGs defined as pHQ under the MIMAG criteria were further screened for the presence of tR-NAs (minimum of 18) and a complete rRNA operon (defined as the presence of at least one copy of each of the 5S, 16S and 23S SSU-rRNA genes, irrespective of whether they were harboured on a single contig or not), and if present were denoted as *high quality* (HQ) MAGs.

### Genome de–replication procedures

We identified putative sets of cognate genomes using the dRep (version 2.2.3) (Olm et al., 2017) `compare` workflow executed with default settings with 20 threads (`-p 20`). Four dereplication analyses were performed; *1*) dereplication of the complete set of MAGs ; *2*) de–replication of the set of 9079 MAGs combining those identified in *1*) with the additional MAGs recovered from using VAMB (**Supplementary Table 7**); *3*) de–replication of the set of 142 HQ MAGs from our analyses combined with the set of 3139 MAGs available from previously published MAG analyses of activated sludge communities (see below) and *4*) The entire set of MAGs from *2*) combined with the 3139 MAGs from references in dRep compare workflow with the same parameters (`-p 80 --S algorithm` `fastANI --multiround primary clustering -sa 0.95 -nc 0.3`) as used in a comparable recent de–replication analysis of rumen MAG catalogues (Watson, 2021).

### Taxonomic and functional annotation of recovered genomes

Taxonomic classification of the collective set of 9079 MAG sequences was performed using the GTDB–Tk (version 0.3.2) (Chaumeil et al., 2019) `classify wf` workflow with default settings (`-x fa --cpus 30`). Prediction of tRNA and rRNA were made using Prokka (version 1.14.6) (Seemann, 2014) executed with default parameters. Predicted rRNA genes were aligned to the SILVA database (version 138.1; release dates 12/06/2020 and 30/06/2020) (Quast et al., 2013) using SINA (version 1.7.1) (Pruesse et al., 2012) with settings `-S --search-min-sim 0.95 -t -v --meta-fmt csv --lca-fields tax slv, tax embl, tax ltp, tax gg, tax rdp`.

### MAG refinement

We performed decontamination of MAG sequences using RefineM (version 0.0.24) (Parks et al., 2017). Briefly, tetranucleotide signature and coverage profiles for contigs were calculated using `scaffold stats` workflow. Contigs with divergent genomic properties were identified with outliers (default settings) and removed using the `filter bins` workflow. Genes were predicted using the `call genes` workflow and annotated with DIA-MOND (Buchfink et al., 2015) against the `gtdb r95 protein db.2020-07-30.dmnd` and `gtdb r95 taxonomy.2020-07-30.tsv` databases, within the `taxon profile` workflow. Contigs with divergent taxonomic assignments were then identified with `taxon filter` and removed with `filter bins`. After decontamination, genome quality was reanalysed using CheckM, as described above, and bins reclassified if indicated.

### Publicly available MAG catalogues

We obtained the following MAG sequence data from the following published studies: 1) a set of 1083 MAGs based on long read metagenome data obtained from 23 wastewater treatment plants in Denmark (Singleton et al., 2021); *2*) a set of 2045 MAGs recovered from meta–analysis of Ye et al. (2020), which includes WWTP samples from several locations in China (data collected by the authors of (Ye et al., 2020)), Singapore (data from (Law et al., 2016)), Denmark (data from (Munck et al., 2015)), USA (data from (Chu et al., 2018)), Argentina (data from (Ibarbalz et al., 2016)), Slovenia (data from (McIlroy et al., 2016)) and Switzerland (data from (Ju et al., 2019)); *3*) one MAG sequence available from NCBI submitted from the time–series metagenome survey of a full–scale activated sludge community in Argentina (Buenos Aires) (Pérez et al., 2019); and *4*) ten MAG sequences available from NCBI from the metagenome survey of three conventional WWTPs in Taiwan inoculated with exogenous anammox pellets (Yang et al., 2020). We re–estimated the genome quality of all MAGs using the CheckM based approach described above.

### Data visualisation

We constructed unrooted phylograms from MAG sequence data using GTDB–Tk (version 0.3.2) based on bacterial single–copy gene sets (bac120 ms gene sets) and imported the `.tree` file into R using the read.tree function from ggtree package (version 2.4.2) (Yu et al., 2017) and subsequently rendered using the ggtree function. Venn diagrams were constructed using the R package VennDiagram (version 1.6.0) (Chen and Boutros, 2011).

## Author Contribution Statement

The study was designed by Y.Y.L, S.W. and R.B.H.W. Y.Y.L and R.B.H.W designed field sampling procedures, which was conducted by L.C.W.L and led by Y.Y.L. L.C.W.L and T.Q.N.N performed gDNA extractions. D.I.D–M and S.C.S advised on gDNA extraction procedures and protocols and generated short read sequencing data. M.A.S.H, K.A. and R.B.H.W designed and performed data analysis. All authors contributed to data interpretation. The manuscript was primarily written by R.B.H.W and M.A.S.H with specific sections contributed by other authors.

## Acknowledgements

This research was supported by the Singapore National Research Foundation and Ministry of Education under the Research Centre of Excellence Programme and by program grants 1102–IRIS–10–02 (S.W., R.B.H.W and S.C.S) from the National Research Foundation (NRF). The computational work was performed in part on resources of the National Supercomputing Centre (NSCC, Singapore) supported by Project 11000984. We thank our numerous colleagues from the Public Utilities Board (Republic of Singapore) for access to facilities and biomass which permitted this work to be undertaken.

## Description of Supplementary Materials

**Supplementary Table 1**

Summary of sequencing read data from each of the 24 samples analysed in this study.

**Supplementary Table 2**

Assembly and binning statistics of each of the 24 individual assemblies and the two co– assemblies (constructed with SPAdes and MetaBAT2)

**Supplementary Table 3**

Summary statistics for the 2,912 secondary (non–redundant) clusters from MAGs recovered by 24 individual assemblies and two co–assemblies (constructed with SPAdes and MetaBAT2)

**Supplementary Table 4**

Summary data for all 7,138 MAGs recovered from 24 individual assemblies and two co– assemblies (constructed with SPAdes and MetaBAT2)

**Supplementary Table 5**

MAGs from 24 individual assemblies and two co–assemblies that possessed completeness of more than 90%

**Supplementary Table 6**

Summary data for the MAGs recovered from VAMB workflow

**Supplementary Table 7**

Secondary (non–redundant) clusters of the complete set of MAGs recovered from all four different workflows (24 individual assemblies, two co-assemblies, and VAMB)

**Supplementary Table 8**

Putative high quality (pHQ) MAGs recovered from all four different workflows

## Notes

https://www.ncbi.nlm.nih.gov/bioproject/PRJNA731554

### Competing Interest Statement

The authors have declared no competing interest.

## References

A. Almeida, S. Nayfach, M. Boland, F. Strozzi, M. Beracochea, Z. J. Shi, K. S. Pollard, E. Sakharova, D. H. Parks, P. Hugenholtz, N. Segata, N. C. Kyrpides, and R. D. Finn. A unified catalog of 204,938 reference genomes from the human gut microbiome. Nature Biotechnology, 39(1):105–114, 2021. ISSN 1546-1696. doi: 10.1038/s41587-020-0603-3. URL https://doi.org/10.1038/s41587-020-0603-3.

I. R. Arkhipova. Metagenome proteins and database contamination. mSphere, 5(6), Nov 2020. ISSN 2379-5042 (Electronic); 2379-5042 (Linking). doi: 10.1128/mSphere.00854-20.

K. Arumugam, C. Bağc, I. Bessarab, S. Beier, B. Buchfink, A. Górska, G. Qiu, D. H. Huson, and R. B. H. Williams. Annotated bacterial chromosomes from frame-shift-corrected long-read metagenomic data. Microbiome, 7(1):61, Apr 2019. ISSN 2049-2618 (Electronic); 2049-2618 (Linking). doi: 10.1186/s40168-019-0665-y.

K. Arumugam, I. Bessarab, M. A. S. Haryono, X. Liu, R. E. Zuniga-Montanez, S. Roy, G. Qiu, D. I. Drautz-Moses, Y. Y. Law, S. Wuertz, F. M. Lauro, D. H. Huson, and R. B. H. Williams. Recovery of complete genomes and non-chromosomal replicons from activated sludge enrichment microbial communities with long read metagenome sequencing. npj Biofilms and Microbiomes, 7(1):23, 2021. doi: 10.1038/s41522-021-00196-6. URL https://doi.org/10.1038/s41522-021-00196-6.

D. Bertrand, J. Shaw, M. Kalathiyappan, A. H. Q. Ng, M. S. Kumar, C. Li, M. Dvornicic, J. P. Soldo, J. Y. Koh, C. Tong, O. T. Ng, T. Barkham, B. Young, K. Marimuthu, K. R. Chng, M. Sikic, and N. Nagarajan. Hybrid metagenomic assembly enables high-resolution analysis of resistance determinants and mobile elements in human microbiomes. Nature Biotechnology, 37(8):937–944, 2019. doi: 10.1038/s41587-019-0191-2. URL https://doi.org/10.1038/s41587-019-0191-2.

D. M. Bickhart, M. Kolmogorov, E. Tseng, D. M. Portik, A. Korobeynikov, I. Tol-stoganov, G. Uritskiy, I. Liachko, S. T. Sullivan, S. B. Shin, A. Zorea, V. P. Andreu, K. Panke-Buisse, M. H. Medema, I. Mizrahi, P. A. Pevzner, and T. P. Smith. Generation of lineage-resolved complete metagenome-assembled genomes by precision phasing. bioRxiv, 2021. doi: 10.1101/2021.05.04.442591. URL https://www.biorxiv.org/content/early/2021/05/04/2021.05.04.442591.

A. Bishara, E. L. Moss, M. Kolmogorov, A. E. Parada, Z. Weng, A. Sidow, A. E. Dekas, S. Batzoglou, and A. S. Bhatt. High-quality genome sequences of uncultured microbes by assembly of read clouds. Nature Biotechnology, 36(11):1067–1075, 2018. doi: 10.1038/nbt.4266. URL https://doi.org/10.1038/nbt.4266.

R. M. Bowers, N. C. Kyrpides, R. Stepanauskas, M. Harmon-Smith, D. Doud, T. B. K. Reddy, F. Schulz, J. Jarett, A. R. Rivers, E. A. Eloe-Fadrosh, S. G. Tringe, N. N. Ivanova, A. Copeland, A. Clum, E. D. Becraft, R. R. Malmstrom, B. Birren, M. Podar, P. Bork, G. M. Weinstock, G. M. Garrity, J. A. Dodsworth, S. Yooseph, G. Sutton, F. O. Glöckner, J. A. Gilbert, W. C. Nelson, S. J. Hallam, S. P. Jungbluth, T. J. G. Ettema, S. Tighe, K. T. Konstantinidis, W.-T. Liu, B. J. Baker, T. Rattei, J. A. Eisen, B. Hedlund, K. D. McMahon, N. Fierer, R. Knight, R. Finn, G. Cochrane, I. Karsch-Mizrachi, G. W. Tyson, C. Rinke, N. C. Kyrpides, L. Schriml, G. M. Garrity, P. Hugenholtz, G. Sutton, P. Yilmaz, F. Meyer, F. O. Glöckner, J. A. Gilbert, R. Knight, R. Finn, G. Cochrane, I. Karsch-Mizrachi, A. Lapidus, F. Meyer, P. Yilmaz, D. H. Parks, A. Murat Eren, L. Schriml, J. F. Banfield, P. Hugenholtz, T. Woyke, and T. G. S. Consortium. Minimum information about a single amplified genome (misag) and a metagenome-assembled genome (mimag) of bacteria and archaea. Nature Biotechnology, 35(8):725–731, 2017. ISSN 1546-1696. doi: 10.1038/nbt.3893. URL https://doi.org/10.1038/nbt.3893.

C. T. Brown, D. Moritz, M. P. O’Brien, F. Reidl, T. Reiter, and B. D. Sullivan. Exploring neighborhoods in large metagenome assembly graphs using spacegraphcats reveals hidden sequence diversity. Genome Biology, 21(1):164, 2020. doi: 10.1186/s13059-020-02066-4. URL https://doi.org/10.1186/s13059-020-02066-4.

B. Buchfink, C. Xie, and D. H. Huson. Fast and sensitive protein alignment using diamond. Nat Methods, 12(1):59–60, Jan 2015. ISSN 1548-7105 (Electronic); 1548-7091 (Linking). doi: 10.1038/nmeth.3176.

R. Cavicchioli, W. J. Ripple, K. N. Timmis, F. Azam, L. R. Bakken, M. Baylis, M. J. Behrenfeld, A. Boetius, P. W. Boyd, A. T. Classen, T. W. Crowther, R. Danovaro, C. M. Foreman, J. Huisman, D. A. Hutchins, J. K. Jansson, D. M. Karl, B. Koskella, D. B. Mark Welch, J. B. H. Martiny, M. A. Moran, V. J. Orphan, D. S. Reay, J. V. Remais, V. I. Rich, B. K. Singh, L. Y. Stein, F. J. Stewart, M. B. Sullivan, M. J. H. van Oppen, S. C. Weaver, E. A. Webb, and N. S. Webster. Scientists’ warning to humanity: microorganisms and climate change. Nat Rev Microbiol, 17(9):569–586, Sep 2019. ISSN 1740-1534 (Electronic); 1740-1526 (Print); 1740-1526 (Linking). doi: 10.1038/s41579-019-0222-5.

P.-A. Chaumeil, A. J. Mussig, P. Hugenholtz, and D. H. Parks. Gtdb-tk: a toolkit to classify genomes with the genome taxonomy database. Bioinformatics, 36(6):1925– 1927, Nov 2019. ISSN 1367-4811 (Electronic); 1367-4803 (Print); 1367-4803 (Linking). doi: 10.1093/bioinformatics/btz848.

H. Chen and P. C. Boutros. Venndiagram: a package for the generation of highly-customizable venn and euler diagrams in r. BMC Bioinformatics, 12(1):35, 2011. doi: 10.1186/1471-2105-12-35. URL https://doi.org/10.1186/1471-2105-12-35.

L.-X. Chen, K. Anantharaman, A. Shaiber, A. M. Eren, and J. F. Banfield. Accurate and complete genomes from metagenomes. Genome Research, 30(3):315–333, 2020. doi: 10.1101/gr.258640.119. URL http://genome.cshlp.org/content/30/3/315.abstract.

B. T. T. Chu, M. L. Petrovich, A. Chaudhary, D. Wright, B. Murphy, G. Wells, and R. Poretsky. Metagenomics reveals the impact of wastewater treatment plants on the dispersal of microorganisms and genes in aquatic sediments. Applied and Environmental Microbiology, 84(5), 2018. ISSN 0099-2240. doi: 10.1128/AEM.02168-17. URL https://aem.asm.org/content/84/5/e02168-17.

M. Z. DeMaere and A. E. Darling. bin3c: exploiting hi-c sequencing data to accurately resolve metagenome-assembled genomes. Genome Biology, 20(1):46, 2019. doi: 10.1186/s13059-019-1643-1. URL https://doi.org/10.1186/s13059-019-1643-1.

J. Ding and A. Regev. Deep generative model embedding of single-cell rna-seq profiles on hyperspheres and hyperbolic spaces. Nat Commun, 12(1):2554, May 2021. ISSN 2041-1723 (Electronic); 2041-1723 (Linking). doi: 10.1038/s41467-021-22851-4.

G. M. Douglas and M. G. I. Langille. Current and Promising Approaches to Identify Horizontal Gene Transfer Events in Metagenomes. Genome Biology and Evolution, 11(10):2750–2766, 08 2019. ISSN 1759-6653. doi: 10.1093/gbe/evz184. URL https://doi.org/10.1093/gbe/evz184.

A. Gurevich, V. Saveliev, N. Vyahhi, and G. Tesler. Quast: quality assessment tool for genome assemblies. Bioinformatics, 29(8):1072–1075, Apr 2013. ISSN 1367-4811 (Electronic); 1367-4803 (Print); 1367-4803 (Linking). doi: 10.1093/bioinformatics/btt086.

R. Hamilton, K. D. Kits, V. A. Ramonovskaya, O. N. Rozova, H. Yurimoto, H. Iguchi, V. N. Khmelenina, Y. Sakai, P. F. Dunfield, M. G. Klotz, C. Knief, H. J. M. Op den Camp, M. S. M. Jetten, F. Bringel, S. Vuilleumier, M. M. Svenning, N. Shapiro, T. Woyke, Y. A. Trotsenko, L. Y. Stein, and M. G. Kalyuzhnaya. Draft genomes of gammaproteobacterial methanotrophs isolated from terrestrial ecosystems. Genome Announc, 3(3), Jun 2015. ISSN 2169-8287 (Print); 2169-8287 (Electronic). doi: 10.1128/genomeA.00515-15.

S. Hofmeyr, R. Egan, E. Georganas, A. C. Copeland, R. Riley, A. Clum, E. EloeFadrosh, S. Roux, E. Goltsman, A. Buluç, D. Rokhsar, L. Oliker, and K. Yelick. Terabase-scale metagenome coassembly with metahipmer. Scientific Reports, 10 (1):10689, 2020. ISSN 2045-2322. doi: 10.1038/s41598-020-67416-5. URL https://doi.org/10.1038/s41598-020-67416-5.

F. M. Ibarbalz, E. Orellana, E. L. M. Figuerola, and L. Erijman. Shotgun metagenomic profiles have a high capacity to discriminate samples of activated sludge according to wastewater type. Applied and Environmental Microbiology, 82(17):5186–5196, 2016. ISSN 0099-2240. doi: 10.1128/AEM.00916-16. URL https://aem.asm.org/content/82/17/5186.

F. Ju, K. Beck, X. Yin, A. Maccagnan, C. S. McArdell, H. P. Singer, D. R. Johnson, T. Zhang, and H. Bürgmann. Wastewater treatment plant resistomes are shaped by bacterial composition, genetic exchange, and upregulated expression in the effluent microbiomes. The ISME Journal, 13(2):346–360, 2019. ISSN 1751-7370. doi: 10.1038/s41396-018-0277-8. URL https://doi.org/10.1038/s41396-018-0277-8.

D. D. Kang, F. Li, E. Kirton, A. Thomas, R. Egan, H. An, and Z. Wang. Metabat 2: an adaptive binning algorithm for robust and efficient genome reconstruction from metagenome assemblies. PeerJ, 7:e7359, 2019. ISSN 2167-8359 (Print); 2167-8359 (Electronic); 2167-8359 (Linking). doi: 10.7717/peerj.7359.

G. A. Kowalchuk and J. R. Stephen. Ammonia-oxidizing bacteria: a model for molecular microbial ecology. Annu Rev Microbiol, 55:485–529, 2001. ISSN 0066-4227 (Print); 0066-4227 (Linking). doi: 10.1146/annurev.micro.55.1.485.

B. Langmead and S. L. Salzberg. Fast gapped-read alignment with bowtie 2. Nat Methods, 9(4):357–359, Mar 2012. ISSN 1548-7105 (Electronic); 1548-7091 (Print); 1548-7091 (Linking). doi: 10.1038/nmeth.1923.

Y. Law, R. H. Kirkegaard, A. A. Cokro, X. Liu, K. Arumugam, C. Xie, M. Stokholm-Bjerregaard, D. I. Drautz-Moses, P. H. Nielsen, S. Wuertz, and R. B. H. Williams. Integrative microbial community analysis reveals full-scale enhanced biological phosphorus removal under tropical conditions. Scientific Reports, 6(1):25719, 2016. ISSN 2045-2322. doi: 10.1038/srep25719. URL https://doi.org/10.1038/srep25719.

H. Li, B. Handsaker, A. Wysoker, T. Fennell, J. Ruan, N. Homer, G. Marth, G. Abecasis, and R. Durbin. The sequence alignment/map format and samtools. Bioinformatics, 25 (16):2078–2079, Aug 2009. ISSN 1367-4811 (Electronic); 1367-4803 (Print); 1367-4803 (Linking). doi: 10.1093/bioinformatics/btp352.

L. M. Lui, T. N. Nielsen, and A. P. Arkin. A method for achieving complete microbial genomes and improving bins from metagenomics data. PLoS Comput Biol, 17(5):e1008972, May 2021. ISSN 1553-7358 (Electronic); 1553-734X (Print); 1553-734X (Linking). doi: 10.1371/journal.pcbi.1008972.

V. Mallawaarachchi, A. Wickramarachchi, and Y. Lin. GraphBin: refined binning of metagenomic contigs using assembly graphs. Bioinformatics, 36(11):3307– 3313, 03 2020. ISSN 1367-4803. doi: 10.1093/bioinformatics/btaa180. URL https://doi.org/10.1093/bioinformatics/btaa180.

M. Martin. Cutadapt removes adapter sequences from high-throughput sequencing reads. EMBnet.journal, 17(1):10–12, 2011. ISSN 2226-6089. doi: 10.14806/ej.17.1.200. URL http://journal.embnet.org/index.php/embnetjournal/article/view/200.

C. Martineau, C. Villeneuve, F. Mauffrey, and R. Villemur. Complete genome sequence of hyphomicrobium nitrativorans strain nl23, a denitrifying bacterium isolated from biofilm of a methanol-fed denitrification system treating seawater at the montreal biodome. Genome Announc, 2(1), Jan 2014. ISSN 2169-8287 (Print); 2169-8287 (Electronic). doi: 10.1128/genomeA.01165-13.

S. J. McIlroy, M. Albertsen, E. K. Andresen, A. M. Saunders, R. Kristiansen, M. Stokholm-Bjerregaard, K. L. Nielsen, and P. H. Nielsen. ‘candidatus competibacter’-lineage genomes retrieved from metagenomes reveal functional metabolic diversity. ISME J, 8(3):613–624, Mar 2014. ISSN 1751-7370 (Electronic); 1751-7362 (Print); 1751-7362 (Linking). doi: 10.1038/ismej.2013.162.

S. J. McIlroy, S. M. Karst, M. Nierychlo, M. S. Dueholm, M. Albertsen, R. H. Kirkegaard, R. J. Seviour, and P. H. Nielsen. Genomic and in situ investigations of the novel uncultured chloroflexi associated with 0092 morphotype filamentous bulking in activated sludge. The ISME Journal, 10(9):2223–2234, 2016. ISSN 1751-7370. doi: 10.1038/is-mej.2016.14. URL https://doi.org/10.1038/ismej.2016.14.

S. L. McLellan, J. C. Fisher, and R. J. Newton. The microbiome of urban waters. Int Microbiol, 18(3):141–149, Sep 2015. ISSN 1139-6709 (Print); 1139-6709 (Linking). doi: 10.2436/20.1501.01.244.

C. Munck, M. Albertsen, A. Telke, M. Ellabaan, P. H. Nielsen, and M. O. A. Sommer. Limited dissemination of the wastewater treatment plant core resistome. Nature Communications, 6(1):8452, 2015. ISSN 2041-1723. doi: 10.1038/ncomms9452. URL https://doi.org/10.1038/ncomms9452.

S. Nayfach, Z. J. Shi, R. Seshadri, K. S. Pollard, and N. C. Kyrpides. New insights from uncultivated genomes of the global human gut microbiome. Nature, 568(7753):505–510, 2019. ISSN 1476-4687. doi: 10.1038/s41586-019-1058-x. URL https://doi.org/10.1038/s41586-019-1058-x.

J. N. Nissen, J. Johansen, R. L. Allesøe, C. K. Sønderby, J. J. A. Armenteros, C. H. Grønbech, L. J. Jensen, H. B. Nielsen, T. N. Petersen, O. Winther, and S. Rasmussen. Improved metagenome binning and assembly using deep variational autoencoders. Nature Biotechnology, 2021. ISSN 1546-1696. doi: 10.1038/s41587-020-00777-4. URL https://doi.org/10.1038/s41587-020-00777-4.

S. Nurk, D. Meleshko, A. Korobeynikov, and P. A. Pevzner. metaspades: a new versatile metagenomic assembler. Genome Res, 27(5):824–834, May 2017. ISSN 1549-5469 (Electronic); 1088-9051 (Print); 1088-9051 (Linking). doi: 10.1101/gr.213959.116.

M. R. Olm, C. T. Brown, B. Brooks, and J.F. Banfield. drep: a tool for fast and accurate genomic comparisons that enables improved genome recovery from metagenomes through de-replication. ISME J, 11(12):2864–2868, Dec 2017. ISSN 1751-7370 (Electronic); 1751-7362 (Print); 1751-7362 (Linking). doi: 10.1038/ismej.2017.126.

A. Orakov, A. Fullam, L. P. Coelho, S. Khedkar, D. Szklarczyk, D. R. Mende, T. S. B. Schmidt, and P. Bork. Gunc: detection of chimerism and contamination in prokaryotic genomes. Genome Biol, 22(1):178, Jun 2021. ISSN 1474-760X (Electronic); 1474-7596 (Print); 1474-7596 (Linking). doi: 10.1186/s13059-021-02393-0.

D. H. Parks, M. Imelfort, C. T. Skennerton, P. Hugenholtz, and G. W. Tyson. CheckM: assessing the quality of microbial genomes recovered from isolates, single cells, and metagenomes. Genome Research, 25:1043–55, 2015.

D. H. Parks, C. Rinke, M. Chuvochina, P.-A. Chaumeil, B. J. Woodcroft, P. N. Evans, P. Hugenholtz, and G. W. Tyson. Recovery of nearly 8,000 metagenome-assembled genomes substantially expands the tree of life. Nature Microbiology, 2 (11):1533–1542, 2017. ISSN 2058-5276. doi: 10.1038/s41564-017-0012-7. URL https://doi.org/10.1038/s41564-017-0012-7.

E. Pasolli, F. Asnicar, S. Manara, M. Zolfo, N. Karcher, F. Armanini, F. Beghini, P. Manghi, A. Tett, P. Ghensi, M. C. Collado, B. L. Rice, C. DuLong, X. C. Morgan, C. D. Golden, C. Quince, C. Huttenhower, and N. Segata. Extensive unexplored human microbiome diversity revealed by over 150,000 genomes from metagenomes spanning age, geography, and lifestyle. Cell, 176(3):649–662.e20, 2019. ISSN 0092-8674. doi: https://doi.org/10.1016/j.cell.2019.01.001. URL https://www.sciencedirect.com/science/article/pii/S0092867419300017.

M. V. Pérez, L. D. Guerrero, E. Orellana, E. L. Figuerola, L. Erijman, and J. McGrath. Time series genome-centric analysis unveils bacterial response to operational disturbance in activated sludge. mSystems, 4(4):e00169–19, Aug. 2019. doi: 10.1128/mSystems.00169-19. URL http://msystems.asm.org/content/4/4/e00169-19.abstract.

E. Pruesse, J. Peplies, and F.O. Glöckner. SINA: Accurate high-throughput multiple sequence alignment of ribosomal RNA genes. Bioinformatics, 28(14):1823–1829, 05 2012. ISSN 1367-4803. doi: 10.1093/bioinformatics/bts252. URL https://doi.org/10.1093/bioinformatics/bts252.

G. Qiu, X. Liu, N. M. M. T. Saw, Y. Law, R. Zuniga-Montanez, S. S. Thi, T. Q. Ngoc Nguyen, P. H. Nielsen, R. B. H. Williams, and S. Wuertz. Metabolic traits of candidatus accumulibacter clade iif strain scelse-1 using amino acids as carbon sources for enhanced biological phosphorus removal. Environ Sci Technol, 54(4):2448–2458, Feb 2020. ISSN 1520-5851 (Electronic); 0013-936X (Linking). doi: 10.1021/acs.est.9b02901.

C. Quast, E. Pruesse, P. Yilmaz, J. Gerken, T. Schweer, P. Yarza, J. Peplies, and F.O. Glöckner. The silva ribosomal rna gene database project: improved data processing and web-based tools. Nucleic Acids Res, 41(Database issue):D590–6, Jan 2013. ISSN 1362-4962 (Electronic); 0305-1048 (Print); 0305-1048 (Linking). doi: 10.1093/nar/gks1219.

C. Quince, T. O. Delmont, S. Raguideau, J. Alneberg, A. E. Darling, G. Collins, and A. M. Eren. Desman: a new tool for de novo extraction of strains from metagenomes. Genome iology, 18:181, Sep 2017a.

C. Quince, A. W. Walker, J. T. Simpson, N. J. Loman, and N. Segata. Shotgun metagenomics, from sampling to analysis. Nature Biotechnology, 35(9):833–844, 2017b. doi: 10.1038/nbt.3935. URL https://doi.org/10.1038/nbt.3935.

C. Quince, S. Nurk, S. Raguideau, R. James, O. S. Soyer, J. K. Summers, A. Limasset, A. M. Eren, R. Chikhi, and A. E. Darling. Metagenomics strain resolution on assembly graphs. bioRxiv, 2020. doi: 10.1101/2020.09.06.284828. URL https://www.biorxiv.org/content/early/2020/09/07/2020.09.06.284828.

R Core Team. R: A Language and Environment for Statistical Computing. R Foundation for Statistical Computing, Vienna, Austria, 2021. URL https://www.R-project.org.

D. Ricaboni, M. Mailhe, S. Khelaifia, D. Raoult, and M. Million. Romboutsia timonensis, a new species isolated from human gut. New Microbes New Infect, 12:6–7, Jul 2016. ISSN 2052-2975 (Print); 2052-2975 (Electronic); 2052-2975 (Linking). doi: 10.1016/j.nmni.2016.04.001.

N. Sangwan, F. Xia, and J. A. Gilbert. Recovering complete and draft population genomes from metagenome datasets. Microbiome, 4(1):8, 2016. doi: 10.1186/s40168-016-0154-5. URL https://doi.org/10.1186/s40168-016-0154-5.

T. Seemann. Prokka: rapid prokaryotic genome annotation. Bioinformatics, 30 (14):2068–2069, Jul 2014. ISSN 1367-4811 (Electronic); 1367-4803 (Linking). doi: 10.1093/bioinformatics/btu153.

C. M. Singleton, F. Petriglieri, J. M. Kristensen, R. H. Kirkegaard, T. Y. Michaelsen, M. H. Andersen, Z. Kondrotaite, S. M. Karst, M. S. Dueholm, P. H. Nielsen, and M. Albertsen. Connecting structure to function with the recovery of over 1000 high-quality metagenome-assembled genomes from activated sludge using long-read sequencing. Nature Communications, 12(1):2009, 2021. ISSN 2041-1723. doi: 10.1038/s41467-021-22203-2. URL https://doi.org/10.1038/s41467-021-22203-2.

C. T. Skennerton, J. J. Barr, F. R. Slater, P. L. Bond, and G. W. Tyson. Expanding our view of genomic diversity in candidatus accumulibacter clades. Environ Microbiol, 17(5):1574–1585, May 2015. ISSN 1462-2920 (Electronic); 1462-2912 (Linking). doi: 10.1111/1462-2920.12582.

R. M. Soo, C. T. Skennerton, Y. Sekiguchi, M. Imelfort, S. J. Paech, P. G. Dennis, J. A. Steen, D. H. Parks, G. W. Tyson, and P. Hugenholtz. An expanded genomic representation of the phylum cyanobacteria. Genome Biol Evol, 6(5):1031–1045, May 2014. ISSN 1759-6653 (Electronic); 1759-6653 (Linking). doi: 10.1093/gbe/evu073.

R. D. Stewart, M. D. Auffret, A. Warr, A. W. Walker, R. Roehe, and M. Watson. Compendium of 4,941 rumen metagenome-assembled genomes for rumen microbiome biology and enzyme discovery. Nature Biotechnology, 37(8):953–961, 2019. doi: 10.1038/s41587-019-0202-3. URL https://doi.org/10.1038/s41587-019-0202-3.

M. Stokholm-Bjerregaard, S. J. McIlroy, M. Nierychlo, S. M. Karst, M. Albertsen, and P. H. Nielsen. A critical assessment of the microorganisms proposed to be important to enhanced biological phosphorus removal in full-scale wastewater treatment systems. Front Microbiol, 8:718, 2017. ISSN 1664-302X (Print); 1664-302X (Electronic); 1664-302X (Linking). doi: 10.3389/fmicb.2017.00718.

B. J. Tully, E. D. Graham, and J. F. Heidelberg. The reconstruction of 2,631 draft metagenome-assembled genomes from the global oceans. Scientific Data, 5(1):170203, 2018. doi: 10.1038/sdata.2017.203. URL https://doi.org/10.1038/sdata.2017.203.

G. V. Uritskiy, J. DiRuggiero, and J. Taylor. Metawrap – a flexible pipeline for genome– resolved metagenomic data analysis. Microbiome, 6(1):158, 2018. doi: 10.1186/s40168-018-0541-1. URL https://doi.org/10.1186/s40168-018-0541-1.

R. Vicedomini, C. Quince, A. E. Darling, and R. Chikhi. Automated strain separation in low-complexity metagenomes using long reads. bioRxiv, 2021. doi: 10.1101/2021.02.24.429166. URL https://www.biorxiv.org/content/early/2021/05/10/2021.02.24.429166.

A. Vijayan, R. K. Vattiringal Jayadradhan, D. Pillai, P. Prasannan Geetha, V. Joseph, and B. S. Isaac Sarojini. Nitrospira as versatile nitrifiers: Taxonomy, ecophysiology, genome characteristics, growth, and metabolic diversity. Journal of Basic Microbiology, 61(2):88–109, 2021. doi: https://doi.org/10.1002/jobm.202000485. URL https://onlinelibrary.wiley.com/doi/abs/10.1002/jobm.202000485.

J. Vollmers, S. Wiegand, and A.-K. Kaster. Comparing and evaluating metagenome assembly tools from a microbiologist’s perspective - not only size matters! PloS One, 12(1):e0169662, 2017. ISSN 1932-6203 (Electronic); 1932-6203 (Linking). doi: 10.1371/journal.pone.0169662.

M. Watson. New insights from 33,813 publicly available metagenome-assembled-genomes (mags) assembled from the rumen microbiome. bioRxiv, 2021. doi: 10.1101/2021.04.02.438222. URL https://www.biorxiv.org/content/early/2021/04/02/2021.04.02.438222.

L. Wu, D. Ning, B. Zhang, Y. Li, P. Zhang, X. Shan, Q. Zhang, M. R. Brown, Z. Li, J. D. Van Nostrand, F. Ling, N. Xiao, Y. Zhang, J. Vierheilig, G. F. Wells, Y. Yang, Y. Deng, Q. Tu, A. Wang, D. Acevedo, M. Agullo-Barcelo, P. J. J. Alvarez, L. Alvarez-Cohen, G. L. Andersen, J. C. de Araujo, K. F. Boehnke, P. Bond, C. B. Bott, P. Bovio, R. K. Brewster, F. Bux, A. Cabezas, L. Cabrol, S. Chen, C. S. Criddle, C. Etchebehere, A. Ford, D. Frigon, J. Sanabria, J. S. Griffin, A. Z. Gu, M. Habagil, L. Hale, S. D. Hardeman, M. Harmon, H. Horn, Z. Hu, S. Jauffur, D. R. Johnson, J. Keller, A. Keucken, S. Kumari, C. D. Leal, L. A. Lebrun, J. Lee, M. Lee, Z. M. P. Lee, M. Li, X. Li, Y. Liu, R. G. Luthy, L.C. Mendonça-Hagler, F. G. R. de Menezes, A. J. Meyers, A. Mohebbi, P. H. Nielsen, A. Oehmen, A. Palmer, P. Parameswaran, J. Park, D. Patsch, V. Reginatto, F. L. de los Reyes, B. E. Rittmann, A. Noyola, S. Rossetti, J. Sidhu, W. T. Sloan, K. Smith, O. V. de Sousa, D. A. Stahl, K. Stephens, R. Tian, J. M. Tiedje, N. B. Tooker, J. D. Van Nostrand, D. De los Cobos Vasconcelos, M. Wagner, S. Wakelin, B. Wang, J. E. Weaver, G. F. Wells, S. West, P. Wilmes, S.-G. Woo, J.-H. Wu, L. Wu, C. Xi, M. Xu, T. Yan, M. Yang, M. Young, H. Yue, T. Zhang, Q. Zhang, W. Zhang, Y. Zhang, H. Zhou, J. Zhou, X. Wen, T. P. Curtis, Q. He, Z. He, P. H. Nielsen, P. J. J. Alvarez, C. S. Criddle, J. M. Tiedje, T. P. Curtis, D. A. Stahl, B. E. Rittmann, and G. W. M. Consortium. Global diversity and biogeography of bacterial communities in wastewater treatment plants. Nature Microbiology, 4(7):1183–1195, 2019. doi: 10.1038/s41564-019-0426-5. URL https://doi.org/10.1038/s41564-019-0426-5.

Y. Yang, J. Pan, Z. Zhou, J. Wu, Y. Liu, J.-G. Lin, Y. Hong, X. Li, M. Li, and J.-D. Gu. Complex microbial nitrogen-cycling networks in three distinct anammox-inoculated wastewater treatment systems. Water Research, 168:115142, 2020. ISSN 0043-1354. doi: 10.1016/j.watres.2019.115142. URL https://www.sciencedirect.com/science/article/pii/S0043135419309169.

L. Ye, R. Mei, W.-T. Liu, H. Ren, and X.-X. Zhang. Machine learning-aided analyses of thousands of draft genomes reveal specific features of activated sludge processes. Microbiome, 8(1):16, 2020. ISSN 2049-2618. doi: 10.1186/s40168-020-0794-3. URL https://doi.org/10.1186/s40168-020-0794-3.

Yu, D. K. Smith, H. Zhu, Y. Guan, and T. T.-Y. Lam. ggtree: an r package for visualization and annotation of phylogenetic trees with their covariates and other associated data. Methods in Ecology and Evolution, 8(1):28–36, 2017. doi: https://doi.org/10.1111/2041-210X.12628. URL https://besjournals.onlinelibrary.wiley.com/doi/abs/10.1111/2041-210X.12628.

